# A unified framework to identify demographic buffering in natural populations

**DOI:** 10.1101/2023.07.03.547528

**Authors:** Gabriel Silva Santos, Samuel J L Gascoigne, André Tavares Corrêa Dias, Maja Kajin, Roberto Salguero-Gómez

## Abstract

The Demographic Buffering Hypothesis (DBH) predicts that natural selection reduces the temporal fluctuations in demographic processes (such as survival, development, and reproduction), due to their negative impacts on population dynamics. However, a comprehensive approach that allows for the examination of demographic buffering patterns across multiple species is still lacking. Here, we propose a three-step framework aimed at identifying and quantifying demographic buffering. Firstly, we categorize species along a continuum of variance based on their stochastic elasticities. Secondly, we examine the linear selection gradients, followed by the examination of nonlinear selection gradients as the third step. With these three steps, our framework overcomes existing limitations of conventional approaches to identify and quantify demographic buffering, allows for multi-species comparisons, and offers an insight into the evolutionary forces that shape demographic buffering. We apply this framework to mammal species and discuss both the advantages and potential of our framework.

---

Environmental stochasticity plays a pivotal role in shaping organisms’ life histories (Bonsall & Klug 2011). Nonetheless, how organisms will cope with the increasing variation in environmental conditions expected under climate change (Boyce *et al*. 2006; Morris *et al*. 2008) is one of the most intriguing questions of ecology and evolution (Sutherland *et al*. 2013).

Evolutionary demography offers a wide array of explanations for the evolutionary processes that shape the diversity of demographic responses to environmental stochasticity (Charlesworth 1994; Pfister 1998; Tuljapurkar *et al*. 2009; Healy *et al*. 2019; Hilde *et al*. 2020). The Demographic Buffering Hypothesis (*DBH*, hereafter) (Morris & Doak 2004; Pélabon *et al*. 2020) predicts a negative relationship between the contribution of a demographic processes (*e.g*., survival, development, reproduction) to the population growth rate (*λ*) and their temporal variance (Pfister 1998)n. The emerging demographic strategy, demographic buffering, accommodates variance of demographic processes to cope with the otherwise negative effects of stochastic environments on *λ* (Pfister 1998; Morris & Doak 2004; Hilde *et al*. 2020).

A unified approach to unambiguously quantify demographic buffering is still missing. Indeed, identifying demographic buffering remains challenging (Morris & Doak 2004; Doak *et al*. 2005) for at least three reasons. First is the different interpretation of results from correlational analyses (*e.g*., as in Pfister, 1998). Some authors have used the correlation coefficient as an index to order species’ life histories in a continuum ranging from buffered (Spearman’s correlation *ρ* = <0 between the sensitivity of *λ* to demographic processes and their temporal variance) to labile (*ρ* = >0, regardless of the “scatterness” around the regression (McDonald *et al*. 2017). In contrast, other researchers interpret the absence of statistical support for demographic buffering as an alternative strategy where variance in demographic process(es) is favoured to track environmental conditions (the so-called Demographic Lability Hypothesis (*DLH*, hereafter; *e.g.,*(Koons *et al*. 2009; Reed & Slade 2012; Jäkäläniemi *et al*. 2013; Hilde *et al*. 2020).

The second obstacle to obtain generalisation across species’ populations regarding demographic buffering is the hierarchical level at which this phenomenon is typically examined. Some studies base their investigations of demographic buffering on the *whole* life history at the level of species or populations (*interspecific level*, hereafter), focusing on the one demographic process that is the most influential for *λ* (Reed & Slade 2012; McDonald *et al*. 2017). At the interspecific level, a life history is referred to as demographically buffered if the most important demographic process has low temporal variance (Pfister 1998; Morris & Doak 2004; Hilde *et al*. 2020; Le Coeur *et al*. 2022). Thus, the associated strategy is commonly decided based on a *single* demographic process (*e.g*., adult survival), ignoring the selection pressures on the rest of the demographic processes within the life cycle. However, to understand how, why, and where demographic buffering occurs –or not– and how buffering patterns might be modified in response to the environment, it is essential to also consider the features *within* a single species’ life cycle (*intraspecific level*, hereafter). Within a single life cycle one demographic process can be buffered against while another can be labile to the environment – supporting the DLH (Koons *et al*. 2009; Jongejans *et al*. 2010; Barraquand & Yoccoz 2013). Thus, for a mechanistic understanding of how environmental stochasticity shapes life histories, both inter-and intra-specific levels need to be addressed.

The third reason limiting a holistic understanding of demographic strategies in stochastic environments are the challenges inherent to examining their underlying mechanisms. Evidence for demographic buffering exists across some long-lived organisms with complex life cycles, (Pfister 1998; Gaillard & Yoccoz 2003; Doak *et al*. 2005; Rotella *et al*. 2012; McDonald *et al*. 2017), but also in short-lived species (Pfister 1998; Reed & Slade 2012; Ferreira *et al*. 2013). Importantly, these patterns of variation do not inform on how the life histories were shaped by natural selection. To do so, one would need to identify the type (linear or nonlinear) and strength of selection acting on demographic processes. Linear selection acts on changing *the mean* value of a demographic process via a linear function between the fitness and the demographic process. In contrast, nonlinear selection acts on *the variance* of demographic processes either increasing it, decreasing it, or increasing/decreasing *the covariance* between two demographic processes (Brodie et al. 1995; Carslake et al. 2008).

The sign of the self-second derivative of *λ* determines the type of nonlinear selection acting on a demographic process. For instance, a negative self-second derivative for a given demographic process describes a concave form of selection, commonly referred to as the ∩-shaped selection (Caswell 1996, 2001; Shyu & Caswell 2014). This form of selection reduces the temporal variance in said demographic process, thereby providing support for the DBH. Conversely, a demographic process yielding a positive self-second derivative identifies a convex, or U-shaped selection (Caswell 1996, 2001; Shyu & Caswell 2014). Such a selection mechanism acts upon demographic processes amplifying their temporal variance, thus supporting the DLH (Koons *et al*. 2009; Le Coeur *et al*. 2022). The cross-second derivatives (not discussed here, see Caswell 1996, 2001 for further details) quantify selection pressures acting on the strength of correlation among different demographic processes.

The rich variation in demographic strategies across the Tree of Life is a result of evolutionary processes that have shaped variance in demographic processes through time. In this context, setting demographic buffering into the adaptive landscape context of linear and nonlinear selection enables us to identify and quantify the evolutionary processes that generate said demographic patterns. In this way, one will better understand how increased variability of environmental conditions might act on the existing –and shape novel– demographic strategies. However, we still lack a unified approach to quantify DBH.

Here, we present a framework that identifies and quantifies demographic buffering. Our framework provides a thorough analysis of temporal variance in demographic processes affected by environmental stochasticity. This framework involves categorizing species or populations along a variance continuum based on the extent to which key demographic processes are buffered by natural selection, thereby limiting their temporal variability. The framework consists of four steps with a mix of well-known methods applied to stage-structured demographic information (*e.g.,* matrix population models [Caswell 2001]; integral projection models [Easterling et al. 2000]). First, we position species or populations on the aforementioned continuum to assess the cumulative effect of the variance on their key demographic processes at the interspecific level (see below). Second, we investigate the presence of linear selection forces operating within the life cycle of each species or population at the intraspecific level (below). Third, we explore the impact of non-linear selection forces acting within the life cycle of each species or population, also at the intraspecific level. The combination of these three steps provides quantitative evidence for/against the DBH, while in step four we describe how to test the DLH.

To demonstrate the applicability of our framework, we apply it to 40 populations of 34 mammal species sourced from the COMADRE database (Salguero-Gómez *et al*. 2016). We showcase how the framework can provide valuable insights into the patterns of demographic buffering across species. The framework offers novel, detailed insights into the selection pressures that act *within* species’ life cycles, thus allowing for a thorough understanding of the evolutionary selection forces that shape the patterns of demographic buffering across species. Beyond providing a quantitative, systematic toolset to test the DBH through three steps, we have also offer an alternative fourth step that briefly outlines how to test for the DLH.

## A unified framework to assess evidence of DBH

The evidence for demographic buffering has been mainly assessed using Matrix Population Models (Pfister 1998; Rotella *et al*. 2012). However, Integral Projection Models (IPM; Rodríguez-Caro et al. 2020; Wang et al. 2023) can be equally applied for identifying the demographic buffering signatures. Both MPMs and IPMs are stage-structured, discrete-time demographic models (Caswell 2001; Ellner *et al*. 2016). For simplicity, here we focus on MPMs, but note that the same approaches are as equally applicable to IPMs (Griffith 2017; Doak *et al*. 2021). Throughout this manuscript, we refer to demographic processes as both matrix entries *aij* (*i.e.,* upper-level parameters) and the vital rates that underline the matrix elements (*i.e.,* lower-level parameters), and note that their conversion is straightforward and described elsewhere (Franco & Silvertown 2004). The framework operates on three steps:

The first step of our framework involves acquiring the relative contribution of each demographic process to the stochastic growth rate, *λ_s_*, the so-called stochastic elasticities, *E*^S^_*ij*_ (Tuljapurkar *et al*. 2003) (Figure 1A). The sum of all stochastic elasticities (*ΣE*^S^_*aij*_), can be separated into two components to assess how temporal variance and mean values of each demographic process contributes to *λ_s_*. The first component represents the *sum of stochastic elasticity of λ_s_ with respect to the variance ΣE*^*S#*^_*aij*_, and the second represents the *sum of stochastic elasticity of λ_s_ with respect to the mean* Σ*λ*^Sμ^, where Σ*λ*^S^ = *ΣE*^*Sσ*^_*aij*_ + Σ*λ*^Sμ^. Thus, the summation *ΣE*^*S#*^_*aij*_ quantifies the extent to which the stochastic population growth rate (*λ_s_*) is influenced by changes in the variances of the demographic processes within the population matrix.

**Figure 1.**
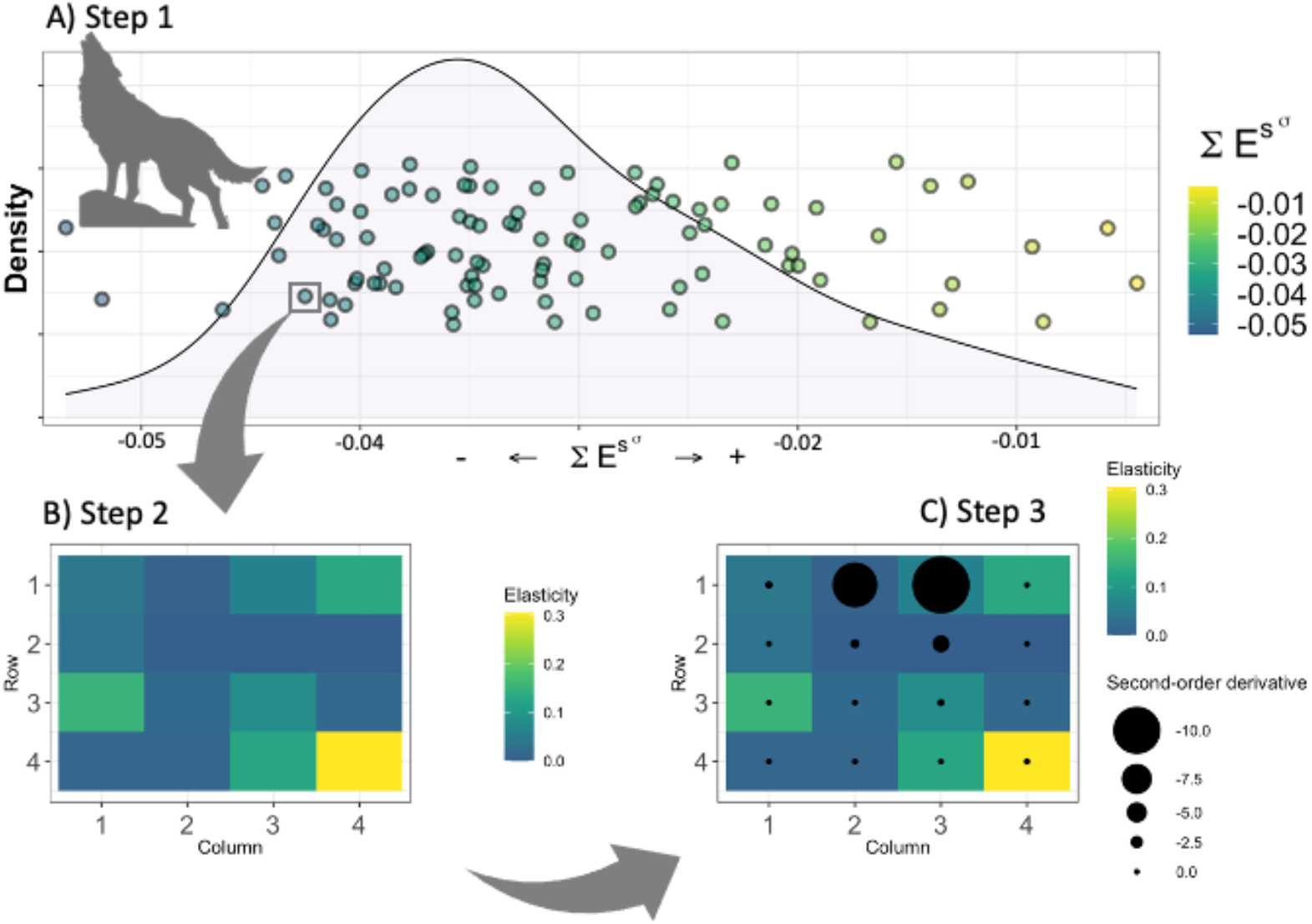
A three-step framework proposed to: Step 1 - allocate species and/or populations on a variance continuum (plot A, dots representing 50 hypothetical species). The variance continuum operates at the interspecific level (see text) and is represented by partitioning the sum of all the stochastic elasticities (*ΣE*^S^_*aij*_) into two compounds: i) sums of stochastic elasticities with respect to the variance (*ΣE*^*Sσ*^_*aij*_), and ii) sums of stochastic elasticities with respect to the mean (Σ*E*^Sμ^). The first step of our framework shows the variance compound of the sums of stochastic elasticities forming a continuum where the right-hand side of the plot represents species (or populations) where a perturbation of variance of the most important demographic process results in weak or no impact on *λ_s_* (yellow dots). The yellow-dotted species (or populations) can be classified as having *buffered life-cycles (supporting the DBH)* – based on the most important demographic process for the *λ_s_*. The left-hand side of the graph represents species (or populations) where a perturbation of the variance of the most important demographic process results in strong impact on *λ_s_* (blue dots). Thus, the blue-dotted species (or populations) can be classified as having *unbuffered life cycles (potentially supporting DLH,* see text) – based on the most important demographic process for the *λ_s_*. The jitter applied on the y-axis has no biological meaning. Step 2 - Access the linear selection pressures for individual species or populations at intraspecific level (see text) (plot B). Step 2 displays the elasticities of the deterministic population growth rate (*λ_t_*) for a hypothetical population of wolf and reveals the linear selection gradients. Step 3 - Access the nonlinear selection pressures at the intraspecific level (see text) (plot C). In the third step self-second derivatives for the corresponding demographic processes from step 2 are displayed.

A higher sum of stochastic elasticity of λ_s_ with respect to the variance (*i.e.,* higher absolute value; |*ΣE*^*Sσ*^_*aij*_|) indicates that small changes in the variance of demographic processes would have a substantial impact on *λ_s_*. In other words, the variance of that demographic process is not constrained by selection, supporting the DLH. On the other hand, a lower (absolute) stochastic elasticity of λ_s_ with respect to the variance suggests that *λ_s_* is less sensitive to such perturbations, or, that variance of such demographic process is being constrained by natural selection, supporting the DBH (Tuljapurkar *et al*. 2003; Haridas & Tuljapurkar 2005) (Fig. 1A).

The first step of the framework thus features the interspecific level and places species or populations alongside a continuum. Species exhibiting unconstrained variance in the most important demographic process (*i.e.,* not buffered/potentially DLH suggesting, Fig. 1A, blue dots) are positioned on the left-hand side of the continuum. In contrast, species with constrained variance in the most important demographic process (*i.e.,* supporting the DBH, Fig. 1A, yellow dots) are positioned on the right-hand side of the continuum. However, the left-hand side of the continuum does not necessarily imply evidence of demographic lability. This is so because demographic lability is defined as an increase in the *mean value* of a demographic process in response to improved environmental conditions (Le Coeur *et al*. 2022). By examining *ΣE*^*Sσ*^_*aij*_, we can visualize an increase or decrease in *variance* of demographic processes, while the mean value of a demographic process does not change. The right-hand side (near 0 values for *ΣE*^*Sσ*^_*aij*_) supports the DBH, while the opposite end represents the lack of support for the DBH, and potentially support for the DLH. However, to undoubtedly provide support for the DLH, further investigation of demographic parameters is needed, as described below.

Step 1 of our framework examines the impacts that environmental variation has on the long-term population growth rate, *λ_s_* (Tuljapurkar *et al*. 2003). This means that the resulting variance continuum in this step of the framework is based on how *λ_s_* was affected by variation in the key demographic parameter across all contiguous time periods.

Steps 2 and 3 of the framework are conducted at the intraspecific level. Once species or populations are positioned along the variance continuum regarding the most important demographic process for *λ_s_*, (step 1), one needs to zoom into each life cycle separately, analysing the selection pressures acting on each one of the demographic processes composing the life cycle. In doing so, one can inspect the selection pressures that have generated the patterns found in step 1. Step 2 (Fig. 1B) requires obtaining the partial derivatives of the deterministic population growth rate, *λ_t_,* relative to all matrix elements of the MPM of interest (*i.e.,* elasticities of *λ_t_* w.r.t each demographic process in the MPM). Step 2 therefore informs on the strength of the natural selection on each of the demographic processes.

Finally, in step 3, one assesses the pattern of nonlinear selection by using the self-second derivatives of *λ_t_* with respect to each demographic process (Fig. 1C). This final step reveals the potential nonlinear selection pressures on all the demographic processes within a life cycle, rather than only the most important one. This final step is key to understanding the evolutionary processes (*i.e.,* types of nonlinear selection) that the demographic processes are subjected to. Without understanding the evolutionary processes operating on the demographic processes, the pattern observed in step 1 might be artefactual. Moreover, step 1 is founded on the assumption that the importance of a demographic process, as indicated by its elasticity, remains unchanged over time. However, stochastic environments can substantially alter elasticity patterns throughout a life cycle (e.g., Lawler et al. 2009).

Steps 2 and 3 of the framework feature selection pressures that have been averaged over the contiguous time periods. This means that the resulting patterns are based on how *λ_t_* (obtained from averaging all sequential MPMs across the duration of the study) would be affected if a demographic process were perturbed. Therefore, steps 2 and 3 are based on a different information than step 1, and can thus complete our understanding of the role of selection pressures on shaping demographic patterns across multiple species.

Another important asset of step 3 above includes the notion that the relative importance (elasticity) of demographic processes themselves changes with changing environment (Stearns 1992). In other words, the extent to which *λ_t_* is sensitive to perturbations in a specific demographic process is *dynamic* (Kroon, Hans *et al*. 2000). Thus, the self-second derivatives generate information on how the sensitivity (or elasticity) of *λ_t_* – based on which the entire variance continuum of species is produced in step 1 – might change. If the sensitivity (or elasticity) of *λ_t_* can change, then it is important to know which demographic processes are most prone to trigger such a change. In the example of a hypothetical wolf species (Fig. 1), this means that if the reproduction of the third age-class individuals (matrix element *a_1,3_*) decreased, the sensitivity of *λ_t_* to *a_1,3_* would increase (square with the largest black dot, Fig. 1C). Consequently, with increased environmental variability, the key demographic process used to place this species onto the variance continuum in step 1 might change from remaining in the fourth age class (matrix element *a_4,4_*, Fig. 1B) to reproduction of the third age-class (matrix element *a_1,3_*, Fig. 1C).

Combining the three steps of our framework allows for the clear, quantitative, holistic identification of evidence to support (or reject) the DBH. Steps 2 and 3 offer key insights as to *why* a given species or population is placed on either the buffered or the non-buffered (potentially labile) end of the variance continuum. A clear and unequivocal evidence for support towards the DBH consists of: (1) a species or population being positioned near the 0 end of the continuum (the right-hand side) in step 1; (2) this species’ or populations’ life cycle having one or more demographic processes with highest elasticity values in step 2; and (3) the same demographic process displaying the highest elasticity in step 2 with negative self-second derivative values in step 3. In this sense, Figure 1B shows that, for the chosen population of a hypothetical wolf species, the most important demographic process is remaining in the fourth stage (MPM element *a_4,4_*), as this demographic process results in highest elasticity value (Fig. 1B yellow square). However, Fig. 1C reveals that *a_4,4_* is under little selection pressure for variance reduction. Thus, there is no evidence for DBH from the third step of the framework (*i.e.,* no concave selection forces), therefore, the lack of concave selection forces on the key demographic process within wolf’s life cycle explains why this species is placed on the left-hand side of the variance continuum (Fig. 1A).

Species placed on the non-buffered end of the continuum is the first but not last step to evidence demographic lability. Indeed, locating a species on the non-buffered end of the variance continuum is a necessary but not sufficient condition for evidence in favour of the DLH. It is key highlighting here that demographic buffering and lability do not represent two extremes of the same continuum. The variance continuum allocates the species or populations from strongly buffered to non-buffered, but to test the DLH, a further step is needed.

Although not our primary goal here, we briefly introduce said step 4. To establish compelling evidence for or against the DLH, it is essential to fulfil several criteria. First, sufficient data across various environments (over time or space) are required to construct reaction norms that depict how a demographic process responds to environmental changes (Morris et al., 2008; Koons et al., 2009). Second, non-linear relationships between demographic processes and the environment must be established based on these reaction norms. Lastly, to identify demographic processes where an increase in the mean value has a stronger positive impact on population growth rate than the detrimental effect of increased variance. This latter condition is only achieved when the vital rate-environment reaction norm is convex (U-shaped; Morris et al. 2008; Koons et al. 2009). Importantly, we note that more likely than previously thought (*e.g.,* Pfister 1998), species do not exist as purely buffering or labile, but that within species, some vital rates may be buffered, other labile, and others insensitive to the environment (*e.g.*, Doak et al. 2005). Deciphering generality in this likely complex pattern should attract much research attention going forward, in our opinion.

## Demographic buffering in mammals: a case study using the unified framework

We demonstrate the performance of our framework using 44 MPMs from 34 mammal species. Mammals are of special interest here for two reasons: (1) mammalian life histories have been well studied (Gillespie 1977; Stearns 1983; Bielby *et al*. 2007; Jones 2011); and (2) some of their populations have already been assessed in terms of buffering, particularly for primates (Morris *et al*. 2008, 2011; Reed & Slade 2012; Rotella *et al*. 2012; Campos *et al*. 2017). Together, the well-studied life histories and previous information about the occurrence of buffering in mammals provide the necessary information to make accurate predictions and validate the performance of the proposed framework.

We used Matrix Population Models from 40 out of 139 studies with mammals available in the COMADRE database v.3.0.0 (Salguero-Gómez *et al*. 2016). These 40 populations encompass 34 species from eight taxonomic orders. We included these MPMs in our analyses because they provide values of demographic processes (*a_ij_*) for three or more contiguous time periods, thus allowing us to obtain the stochastic elasticity of each *a_ij_*.

Although we are aware that not all possible temporal variation in demographic processes may have been expressed within this period, we assumed three or more transitions are enough to provide sufficient variation for population comparison. At least three contiguous time periods - a common selection criteria in comparative studies of stochastic demography (Compagnoni *et al*. 2023) - also allowed to test and showcase our framework. Fortunately, several long-lived species, characterized by low variation in their demographic processes, were studied for a long time (*e.g.,* some primates in our dataset have been studied for over 20 years – Morris *et al*. 2011). We removed the populations where either only survival or only reproduction rates were reported, because of the impossibility to calculate the stochastic growth rate. A detailed description of the analysed data and their original sources are available in supplementary material (Supplementary Material, Table S1).

*Homo sapiens* was included in our analyses because it is the only mammalian species in which second-order derivatives have been applied (Caswell 1996). Therefore, *Homo sapiens* provides an ideal basis for comparisons among species. The data for *Homo sapiens* were gathered from 26 modern populations located in various cities, allowing us to construct a spatiotemporal variance. It is important to note that in this case, we are not working with true temporal variance but rather a variance that encompasses both spatial and temporal aspects.

For steps 2 and 3 of our framework, we utilized a subset of 16 populations (including *Homo sapiens*) whose population projection matrices (MPMs) were organized by age. We specifically selected these populations because their life cycles can be summarized by two main demographic processes: survival and contribution to recruitment of new individuals.

The contribution to recruitment can be interpreted as either the mean reproductive output for each age class or an approximation thereof, depending on how the matrices are structured (Ebert 1999). One advantage of using such matrices is that they encompass only two types of demographic processes, namely survival and recruitment, eliminating the need to account for multiple transitions between different life stages.

To perform the step 1 of our framework and obtain the *ΣE*^*S*^_*aij*_ (and Σ*E*^Sμ^), we followed Tuljapurkar *et al*. (2003). To perform step 2 of our framework, we calculated the deterministic elasticities of each demographic process extracted using the *popbio* package. All analyses were performed using R version 3.5.1 (R Core team, 2018). Finally, to perform the step 3 of our framework the self-second derivatives were adapted from *demogR* (Jones 2007) following Caswell 1996 and applied for the mean MPM.

## Results

We ranked 40 populations from the 34 identified mammal species according to the cumulative impact of variation in demographic processes on *λ_s_* using the step 1 of our framework (Fig. 2). Additional information is provided in the supplementary material (Table S1). Most of the analysed orders were placed on the low-variance end of the variance continuum (Fig. 2). The smallest contributions of variation in demographic processes (*i.e.,* maximum value of *ΣE*^*S*^_*aij*_, note that *ΣE*^*S*^_*aij*_ ranges from 0 to -1), suggesting more buffered populations, were assigned to Primates: northern muriqui (*Brachyteles hyphoxantus*, *ΣE*^*S*^_*aij*_ = - 0.09 × 10^-4^ ± 0.12 × 10^-4^) (mean ± standard deviation) (Fig. 2 silhouette a), mountain gorilla (*Gorilla beringhei*, *ΣE*^*S*^_*aij*_ = -0.24 × 10^-4^ ± 0.08 × 10^-4^) (Fig. 2 silhouette b), followed by the blue monkey (*Cercopithecus mitis*, *ΣE*^*S*^_*aij*_ = -0.63 × 10^-4^ ± 0.06 × 10^-4^) (Fig. 2 silhouette c).

**Figure 2.**
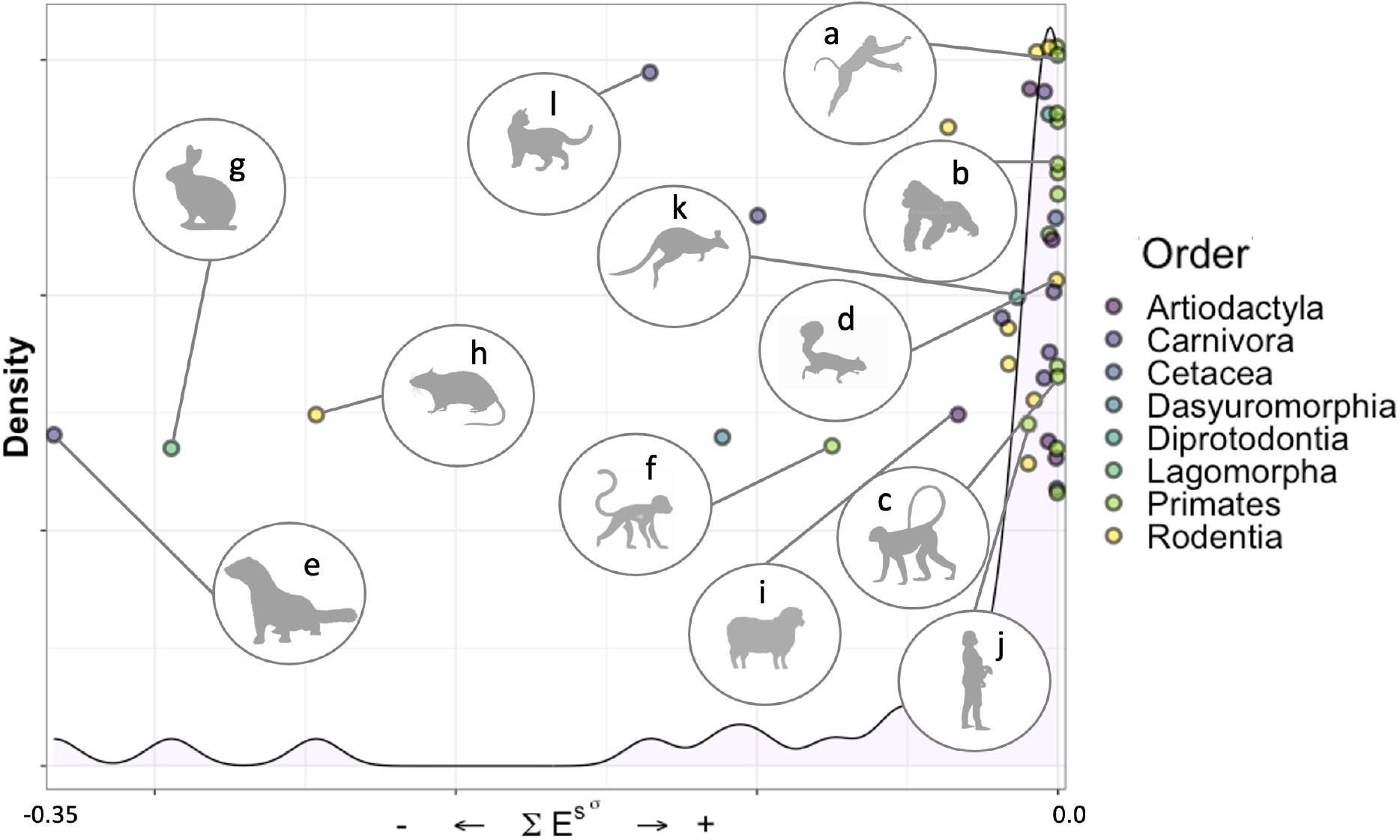
Results for step 1 of our framework showing the sum of stochastic elasticities with respect to the variance *ΣE*^*Sσ*^_*aij*_ increase caused by a perturbation in the most relevant demographic process. The 40 populations from 34 species of mammals from the COMADRE database are ranked into the variance continuum from strongly buffered (right-hand side, supporting the DBH) to more variable, less buffered (left-hand side, potentially supporting the DLH, see text). Colors represent different taxonomic orders with Primates occupying the right-hand side. Silhouettes: a) *Brachyteles hyphoxantus*, b) *Gorilla beringhei*, c) *Cercopithecus mitis*, d) *Urocitellus columbianus,* e) *Mustela erminea*, f) *Erythrocebus patas*, g) *Lepus americanus*, h) *Rattus fuscipes*, i) *Ovis aries*, j) *Homo sapiens*, k) *Macropus eugenii*, and l) *Felis catus*. The jitter applied on the y-axis has no biological meaning.

The first non-primate species placed near the low-variance end of the continuum was the Columbian ground squirrel (*Urocitellus columbianus*, Rodentia, *ΣE*^*S*^_*aij*_ = -0.003 ± 0.002) (Fig. 2 silhouette d). The species with the highest contribution of variation in demographic processes placed at the high-variance end of the continuum was the stoat (*Mustela erminea*, Carnivora, *ΣE*^*S*^_*aij*_ = -0.35 ± 0.02) (Fig. 2 silhouette e). All the 14 primate populations supported the DBH, occupying the right-hand side of the variance continuum, with exception of the Patas monkey (*Erythrocebus patas*, Primates, *ΣE*^*S*^_*aij*_ = -0.05 ± 0.03) (Fig. 2 silhouette f). The snowshoe hare (*Lepus americanus*, Lagomorpha, *ΣE*^*S*^_*aij*_ = -0.29 ± 0.16) (Fig. 2 silhouette g) and the Bush rat (*Rattus fuscipes*, Rodentia, *ΣE*^*S*^_*aij*_ = -0.25 ± 0.03) (Fig. 2 silhouette h) appear on the high-variance end of the continuum.

As predicted for the steps 2 and 3, we could not observe a clear pattern in support of the DBH. This finding means that the demographic processes with the highest elasticity values failed to display strongly negative self-second derivatives (Fig. 3). Particularly for majority of primates - with the lack or minor temporal variation in demographic processes - demographic processes with high elasticities had positive values for the self-second derivatives (indicated by yellow squares with white dots in Fig. 3). Examples of primate species exhibiting high elasticities and positive values for the self-second derivatives and include northern muriqui (*Brachyteles hypoxanthus*), mountain gorilla (*Gorilla beringei*), white-faced capuchin monkey (*Cebus capucinus*), rhesus monkey (*Macaca mulatta*), blue monkey (*Cercopithecus mitis*), Verreaux’s sifaka (*Propithecus verreauxi*) and olive baboon (*Papio cynocephalus*) (Fig. 3). This implies that the key demographic processes influencing *λ_t_* are not subject to selective pressure for reducing their temporal variability. However, even though the primates were positioned closer to the low-variance end of the continuum in step 1, the evidence from steps 2 and 3 does not support DBH.

**Figure 3:**
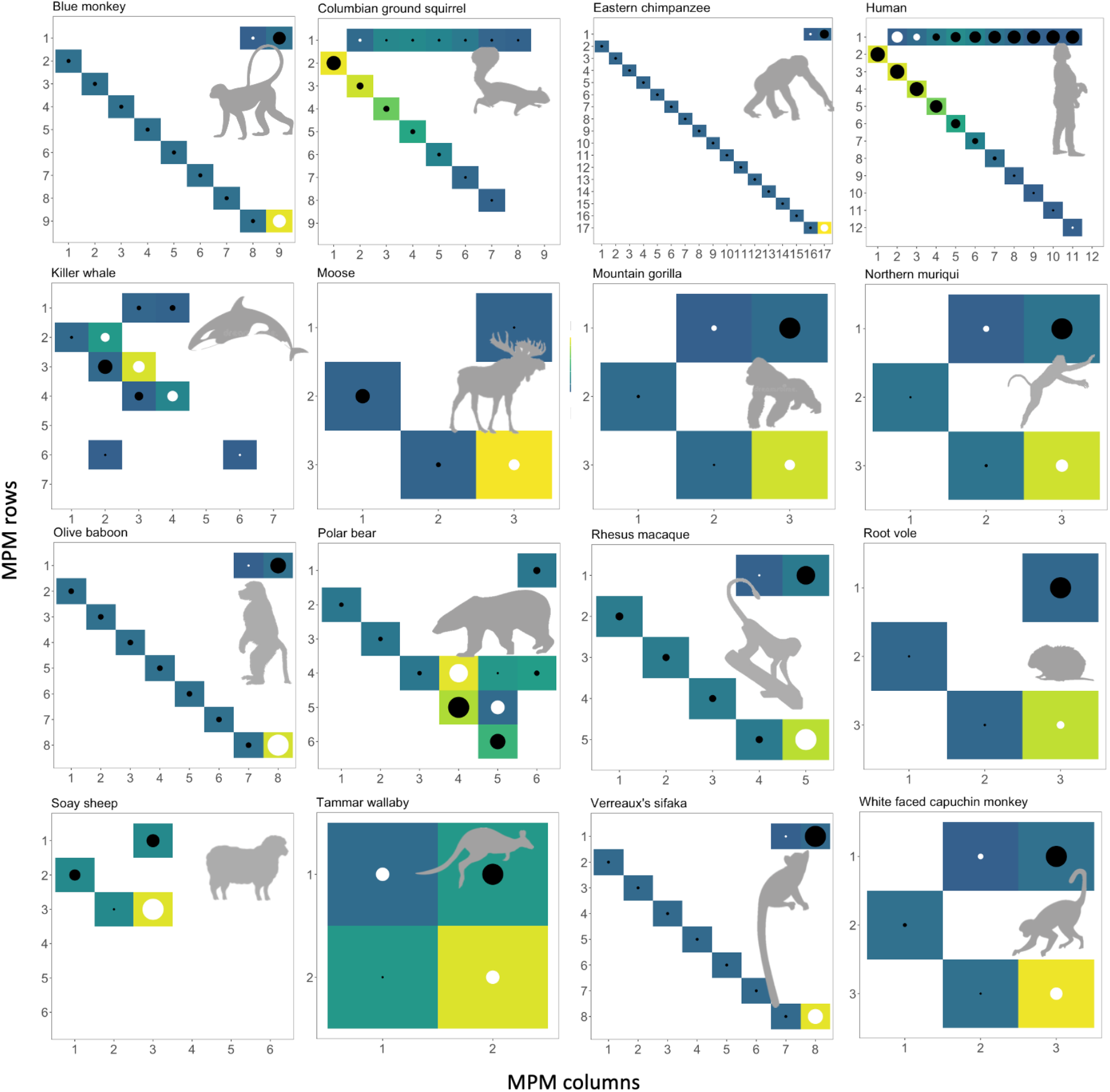
Results from steps 2 and 3 of the proposed framework (see Fig. 2B, C). The 16 plots represent populations where the MPMs built by ages were available in the COMADRE database (see text). The color scale represents elasticity values for each of the demographic processes in the MPM, where yellow represents high and blue low elasticity values. No color means elasticity=0. Because the aim of step 2 is to identify the most important demographic process within each species’ life cycle (the intraspecific level, see text) - not to compare the elasticity values among species - each plot has its own scale (see end of legend). The black dots represent negative self-second derivatives of *λ_t_* - thus concave selection - and the white dots represent positive self-second derivatives of *λ_t_* - thus convex selection. The dot sizes are scaled by the absolute value of self-second derivatives, where the smaller the dot, the closer a self-second derivative is to 0, indicting weak or no selection. Large dots indicate strong selection forces. Scales (E_min-max_=elasticity minimum and maximum value, SSD_min-max_=self-second derivative minimum and maximum value): Blue monkey E_min-max_=0.00-0.52, SSD_min-max_=-1.25-1.27; Columbian ground squirrel: E_min-max_=0.00-0.23, SSD_min-max_=-1.48-0.01; Eastern chimpanzee: E_min-max_=0.00-0.60, SSD_min-max_=-4.39-2.59; Human: E_min-max_=0.00-0.18, SSD_min-max_=-0.15-0.08; Killer whale: E_min-max_=0.00-0.55, SSD_min-max_=-5.72-3.43; Moose: E_min-max_=0.00-0.55, SSD_min-max_=-0.66-0.36; Mountain gorilla: E_min-max_=0.00-0.81, SSD_min-max_=-1.46-0.28; Northern muriqui: E_min-max_=0.00-0.72, SSD_min-max_=-1.17-0.35; Olive baboon: E_min-max_=0.00-0.54, SSD_min-max_=-0.57-1.13; Polar bear: E_min-max_=0.00-0.26, SSD_min-max_=-0.73-0.54; Rhesus macaque: E_min-max_=0.00-0.51, SSD_min-max_=-0.54-0.71; Root vole: E_min-max_=0.00-0.86, SSD_min-max_=-2.54-0.22; Soay sheep: E_min-max_=0.00-0.56, SSD_min-max_=-0.22-0.40; Tammar wallaby: E_min-max_=0.00-0.55, SSD_min-max_=-0.64-0.34; White faced capuchin monkey: E_min-max_=0.00-0.66, SSD_min-max_=-2.66-1.21.

The killer whale showed similar controversy between step 1 and steps 2-3 results as most primates. In step 1, the killer whale was positioned at the buffered end of the variance continuum (*Orcinus orca*, Cetacea, *ΣE*^*S*^_*aij*_ = -0.70 × 10^-4^ ± 1.04 × 10^-5^) (Fig. 2 silhouette not shown). However, steps 2 and 3 show that the three demographic processes in killer whale life cycle with highest elasticity values (matrix elements *a_2,2_*, *a_3,3_* and *a_4,4_*) are not under selection pressures for reducing their temporal variance, but the opposite (depicted by yellow and green squares with white dots, Fig. 3).

The only primate species exhibiting DBH evidence in steps 2 and 3 was human. In human, demographic parameters representing survival from first to second age class (matrix element *a_2,1_*) displayed high elasticities and negative self-second derivatives (depicted as yellow squares with black dots in Fig. 3). Evidence supporting the DBH was also found in the Columbian ground squirrel (*Urocitellus columbianus*), where, similar to humans, survival from the first to the second age class (matrix element *a_2,1_*) showed indications of selection acting to reduce its variance. Accordingly, the Columbian ground squirrel was positioned close to the buffered end of the variance continuum in step 1. Hence, the Columbian ground squirrel was the sole species with consistent DBH support across all three steps of the framework.

The Soay sheep (*Ovis aries*) was the species furthest from the buffered end of the variance continuum that enabled to perform steps 2 and 3. For the Soay sheep, remaining in the third age class (matrix element *a_3,3_*) has the major influence on *λ_t_* and is under selection pressure to have its variance increased. The latter characteristics reveal conditions for the DLH support even though the species is placed closer to the buffered end of the variance continuum.

Steps 2 and 3 illustrate the importance of examining DBH evidence on the intraspecific level. These two steps of the framework identify the simultaneous acting of concave and convex selection on different demographic processes but within a single life cycle. In polar bear (*Ursus maritimus*), the key demographic process (matrix element *a_4,4_*) is under convex selection, as depicted by a yellow square with a white dot in Fig. 3. However, the demographic process with the second highest elasticity value (matrix element *a_5,4_*) is under strong concave selection (depicted by a light green square with a black dot in Fig. 3).

By adding step 3 to the framework, another important information was added. The high absolute values of self-second derivatives (large dots, either black or white, Fig. 3) indicate where the sensitivity of *λ_t_* to demographic parameters is itself prone to environmental changes. For instance, if the value of *a_5,4_* for polar bear increased, the sensitivity of *λ_t_* to *a_5,4_* would decrease because the self-second derivative of *a_5,4_* is highly negative (depicted by the largest black dot in polar bear MPM). Vice versa holds for the *a_4,4_* demographic process, where an increase in the value of *a_4,4_* would increase *λ_t_*’s sensitivity to *a_4,4_*, because the self-second derivative of *a_5,4_* is highly positive (depicted by the largest white dot in polar bear MPM). Thus, sensitivities (or equally elasticities) of demographic processes with high absolute values for self-second derivatives can easily change - potentially changing the key demographic process used for allocating species into the variance continuum in step 1 of the framework.

## Discussion

In the Anthropocene, identifying and quantifying mechanisms of species responses to stochastic environments holds crucial importance. This importance is particularly tangible in the context of the unprecedented environmental changes and uncertainties that impact the dynamics and persistence of natural populations (Boyce *et al*. 2006). Correlational demographic analysis, whereby the importance of demographic processes and their temporal variability is examined (e.g., Pfister 1998), has attempted to identify how species may buffer against the negative effects of environmental stochasticity. However, these widely used approaches have important limitations (see Introduction and Hilde et al. 2020). Our novel framework overcomes said limitations by providing a rigorous approach to test the demographic buffering hypothesis (DBH; Pfister 1998; Hilde et al. 2020).

Evidencing demographic buffering is not straightforward. Indeed, through the analysis of stochastic population growth rate (*λ_s_*) in our application of the framework to 44 populations of 34 species, we identify the highest density of natural populations near the buffered end of the variance continuum (step 1), indicating possible support for the DBH. However, we show that the same species then fail to exhibit signs of concave (∩-shaped) selection on the key demographic parameters when further analyses are performed averaging the variation across the duration of each study (steps 2 and 3). This finding confirms that placing the species near the buffered end of the variance continuum is *necessary* but not *sufficient* to test the DBH. Indeed, buffering occurs when concave selection forces act on the key demographic parameter (Caswell 1996, 2001; Shyu & Caswell 2014).

Combining the three steps into a unified framework is of outmost importance. In steps 2 and 3 of the framework, we find relatively limited overall support for the DBH in the examination of our 16 (out of 34 in step 1) studied animal species. Step 3 of our framework reveals that the role of natural selection shaping temporal variation in demographic processes is more complex than expected by the DBH alone. Indeed, demographic processes within our study populations are often under a mix of convex and concave selection. This mix of selection patterns was already suggested by Doak *et al*. (2005). Here, only two out of 16 mammal species revealed concave selection acting on the key demographic processes (Columbian ground squirrel [*Urocitellus columbianus*], and humans, [*Homo sapiens sapiens*]). These two species were also placed near the buffered end of the variance continuum, therefore meeting all the necessary conditions to diagnose clear support in favour of DBH. However, finding 12.5% (two out of 16) species that meet the criteria for demographic buffering is not in concordance with previous studies. Support for the DBH has been reported across 22 ungulate species (Gaillard & Yoccoz 2003). In the one ungulate we examined, the moose (*Alces alces*), we find only partial support for DBH in adult survival, since this species is placed near the buffered end of the variance continuum in step 1 but does not show concave selection pressures on adult survival in step 2/3, as predicted by the DBH.

Our overall findings reveal varying levels of support for the notion that adult survival in long-lived species tends to be buffered. Indeed, Gaillard *et al*. (1998) found that adult female survival varied considerably less than juvenile survival in large herbivores. This finding was also supported by further studies in ungulates (Gaillard & Yoccoz 2003), turtles (Heppell 1998), vertebrates and plants (Pfister 1998), and more recently across nine (out of 73) species of plants (McDonald *et al*. 2017).

When placing our study species along a variance continuum (step 1), primates tend to be located on the buffered end. However, most primates displayed convex –instead of the expected concave– selection on adult survival. Similar results, where the key demographic process failed to display constrained temporal variability, have been reported for long-lived seabirds (Doherty *et al*. 2004). One explanation for the unexpected convex selection on adult survival involves trade-offs, as suggested by Doak et al. (2005). When two demographic parameters are negatively correlated, the variance of population growth rate (*λ*) can be increased or decreased (Evans & Holsinger 2012; Compagnoni *et al*. 2016). The well-established trade-off between survival and fecundity (e.g., Stearns 1992; Roff & Fairbairn 2007) might explain the observed concave selection signatures on late fecundity and convex selection on adult survival. Because variation in primate recruitment is already constrained by physiological limitations (Campos *et al*. 2017), when adult survival and recruitment are engaged in a trade-off, this trade-off might lead to our unexpected result. Here, future studies may benefit from deeper insights via cross-second derivatives (Caswell 1996, 2001) to investigate correlations among demographic processes.

Examining the drivers of demographic buffering has become an important piece of the ecological and evolutionary puzzle of demography. As such, testing the DBH can help us better predict population responses to environmental variability, climate change, and direct anthropogenic disturbances (Pfister 1998; Boyce *et al*. 2006; McDonald *et al*. 2017; Vázquez *et al*. 2017). By setting the DBH into a broader and integrated framework, we hope to enhance comprehension and prediction of the implications of heightened environmental stochasticity on the evolution of life history traits. This understanding is crucial in mitigating the risk of extinction for the most vulnerable species.

## Acknowledgements

This study was financed in part by the *Coordenação de Aperfeiçoamento de Pessoal de Nível Superior* - Brasil (CAPES) - Finance Code 001. GSS was supported by CAPES and CNPq (301343/2023-3). RS-G was supported by a NERC Independent Research Fellowship (NE/M018458/1). MK was supported by the European Commission through the Marie Skłodowska-Curie fellowship (MSCA MaxPersist #101032484) hosted by RSG.

## Data availability

The demographic data used in this paper are open-access and available in the COMPADRE Plant Matrix Database (v. 5.0.1; https://compadre-db.org/Data/ Compadre). A list of the studies and species used here is available in Supplementary Material (Table S1).

## Supplementary material – Data available in COMADRE Version 2.0.1 and results from Step 1 of the framework

**Table S1.** The metadata used in step 1 of our framework and the respective results presented in the main text. The first four columns represent the information from where Matrix Populations Models (MPMs) were extract precisely as presented in COMADRE 2.0.1. Column titles differ from the database as “SpeciesAuthorComadre” is equivalent to “SpeciesAuthor” and “SpeciesName” is equivalent to “SpeciesAccepted” in COMADRE 2.0.1. The remaining columns present the results of step 1, where we present the raw values of Σ*E*^Sμ^ and *ΣE*^*S*^_*aij*_, their respective standard deviation, the stochastic population growth rate *λ_s_*, and the number of available matrices (# matrices). For ByAge, “TRUE” was assigned for MPMs built by age or “FALSE” if otherwise.

| SpeciesAuthorComadre | SpeciesName | CommonName | Order | $\Sigma E_{a_{ij}}^{S^{\mu}}$ | $\Sigma E_{a_{ij}}^{S^{\mu}}$ (sd) | $\Sigma E_{a_{ij}}^{S^{\sigma}}$ | $\Sigma E_{a_{ij}}^{S^{\sigma}}$ (sd) | # matrices | $\lambda$ |
| --- | --- | --- | --- | --- | --- | --- | --- | --- | --- |
| Homo_sapiens_subsp._sapiens | *Homo sapiens sapiens* | Human | Primates | 1.003 | 0.003 | 1.003 | 0.004 | 13 | 1.064 |
| Alces_alces | *Alces alces* | Moose | Artiodactyla | 1.001 | 0.001 | 1.001 | 0.001 | 13 | 1.205 |
| Antechinus_agilis | *Antechinus agilis* | Agile antechinus | Dasyuromorphia | 1.111 | 0.111 | 1.111 | 0.011 | 2 | 0.931 |
| Brachyteles_hypoxanthus | *Brachyteles hypoxanthus* | Northern muriqui | Primates | 1.000 | 0.000 | 1.000 | 0.000 | 12 | 1.051 |
| Callospermophilus_lateralis | *Callospermophilus lateralis* | Golden-mantled ground squirrel | Rodentia | 1.054 | 0.054 | 1.054 | 0.055 | 9 | 2.052 |
| Cebus_capucinus | *Cebus capucinus* | White faced capuchin monkey | Primates | 1.000 | 0.000 | 1.000 | 0.000 | 11 | 1.021 |
| Cercopithecus_mitis | *Cercopithecus mitis* | Blue monkey | Primates | 1.000 | 0.000 | 1.000 | 0.000 | 14 | 1.036 |
| Eumetopias_jubatus | *Eumetopias jubatus* | Northern sea lion; Steller sea lion | Carnivora | 1.005 | 0.005 | 1.005 | 0.002 | 2 | 0.904 |
| Felis_catus | *Felis catus* | Feral cat | Carnivora | 1.136 | 0.136 | 1.136 | 0.012 | 1 | 1.948 |
| Gorilla_beringei | *Gorilla beringei* | Mountain gorilla | Primates | 1.000 | 0.000 | 1.000 | 0.000 | 21 | 1.027 |
| Hippocamelus_bisulcus | *Hippocamelus bisulcus* | Huemul deer | Artiodactyla | 1.002 | 0.002 | 1.002 | 0.001 | 1 | 0.996 |
| Lepus_americanus | *Lepus americanus* | Snowshoe hare | Lagomorpha | 1.294 | 0.294 | 1.294 | 0.165 | 2 | 0.812 |
| Lycaon_pictus | *Lycaon pictus* | African wild dog | Carnivora | 1.100 | 0.100 | 1.100 | 0.008 | 1 | 1.500 |
| Macaca_mulatta_3 | *Macaca mulatta* | Rhesus macaque | Primates | 1.000 | 0.000 | 1.000 | 0.001 | 12 | 1.127 |
| Macropus_eugenii | *Macropus eugenii* | Tammar wallaby | Diprotodontia | 1.013 | 0.013 | 1.013 | 0.012 | 7 | 0.981 |
| Marmota_flaviventris_2 | *Marmota flaviventris* | Yellow-bellied marmot | Rodentia | 1.007 | 0.007 | 1.007 | 0.006 | 4 | 0.890 |
| Marmota_flaviventris_3 | *Marmota flaviventris* | Yellow-bellied marmot | Rodentia | 1.008 | 0.008 | 1.008 | 0.005 | 4 | 0.921 |
| Microtus_oecconomus | *Microtus oecconomus* | Root vole | Rodentia | 1.000 | 0.000 | 1.000 | 0.001 | 14 | 1.028 |
| Mustela_erminea | *Mustela erminea* | Stoat | Carnivora | 1.334 | 0.334 | 1.334 | 0.117 | 2 | 1.258 |
| Orcinus_orca_2 | *Orcinus orca* | Killer whale | Cetacea | 1.001 | 0.001 | 1.001 | 0.001 | 24 | 0.999 |
| Ovis_aries_2 | *Ovis aries* | Soay sheep | Artiodactyla | 1.033 | 0.033 | 1.033 | 0.020 | 3 | 1.099 |
| Pan_troglodytes_subsp._schweinfurthii | *Pan troglodytes* | Eastern chimpanzee | Primates | 1.000 | 0.000 | 1.000 | 0.001 | 22 | 0.982 |
| Papio_cynocephalus | *Papio cynocephalus* | Olive baboon | Primates | 1.000 | 0.000 | 1.000 | 0.000 | 19 | 1.054 |
| Peromyscus_maniculatus_2 | *Peromyscus maniculatus* | Deer mouse | Rodentia | 1.010 | 0.010 | 1.010 | 0.005 | 2 | 1.107 |
| Phocarcos_hookeri | *Phocarcos hookeri* | New Zealand sea lion | Carnivora | 1.005 | 0.005 | 1.005 | 0.003 | 8 | 1.023 |
| Propithecus_verreauxi | *Propithecus verreauxi* | Verreaux's sifaka | Primates | 1.000 | 0.000 | 1.000 | 0.000 | 12 | 0.986 |
| Puma_concolor_8 | *Puma concolor* | Cougar | Carnivora | NA | NA | NA | NA | 10 | 1.115 |
| Rattus_fuscipes | *Rattus fuscipes* | Bush rat | Rodentia | 1.246 | 0.246 | 1.246 | 0.029 | 2 | 1.305 |
| Spermophilus_armatus | *Urocitellus armatus* | Uinta ground squirrel | Rodentia | 1.016 | 0.016 | 1.016 | 0.011 | 4 | 1.125 |
| Spermophilus_armatus_2 | *Urocitellus armatus* | Uinta ground squirrel | Rodentia | 1.017 | 0.017 | 1.017 | 0.010 | 3 | 1.095 |
| Spermophilus_columbianus | *Urocitellus columbianus* | Columbian ground squirrel | Rodentia | 1.036 | 0.036 | 1.036 | 0.025 | 3 | 1.009 |
| Spermophilus_columbianus_3 | *Urocitellus columbianus* | Columbian ground squirrel | Rodentia | 1.003 | 0.003 | 1.003 | 0.006 | 3 | 1.200 |
| Ursus_americanus_subsp._floridanus | *Ursus americanus* | Florida black bear | Carnivora | 1.003 | 0.003 | 1.003 | 0.003 | 2 | 1.020 |
| Ursus_arctos_subsp._horribilis_5 | *Ursus arctos* | Grizzly bear | Carnivora | 1.001 | 0.001 | 1.001 | 0.001 | 4 | 1.026 |
| Ursus_maritimus_2 | *Ursus maritimus* | Polar bear | Carnivora | 1.019 | 0.019 | 1.019 | 0.007 | 2 | 0.941 |
| Brachyteles_hypoxanthus_2 | *Brachyteles hypoxanthus* | Northern muriqui | Primates | 1.000 | 0.000 | 1.000 | 0.000 | 12 | 1.111 |
| Cebus_capucinus_2 | *Cebus capucinus* | WhiteNAfaced capuchin monkey | Primates | 1.000 | 0.000 | 1.000 | 0.000 | 11 | 1.059 |
| Chlorocebus_aethiops_2 | *Chlorocebus aethiops* | Vervet | Primates | 1.075 | 0.075 | 1.075 | 0.087 | 5 | 1.187 |
| Erythrocebus_patas | *Erythrocebus patas* | Patas monkey | Primates | 1.051 | 0.051 | 1.051 | 0.038 | 5 | 1.128 |
| Gorilla_beringei_subsp._beringei | *Gorilla beringei* | Mountain gorilla | Primates | 1.000 | 0.000 | 1.000 | 0.000 | 21 | 1.053 |

